# Visual occipito-temporal sensitivity to digits through elementary school

**DOI:** 10.1101/2021.10.22.465440

**Authors:** Gorka Fraga-González, Sarah V. Di Pietro, Georgette Pleisch, Jasmin Neuenschwander, Susanne Walitza, Daniel Brandeis, Iliana I. Karipidis, Silvia Brem

## Abstract

Number processing abilities are important for academic and personal development. The course of initial specialization of ventral occipito-temporal cortex (vOTC) for visual number processing is crucial for the development of numeric and arithmetic skills. We examined the visual N1, the electrophysiological correlate of vOTC activations across five time points in kindergarten (T1), middle and end of first grade (T2, T3), second (T4) and fifth grade (T5). 62 children (35 female) performed a target detection task which included visual presentation of digits, false fonts, and letters. Arithmetic skills were measured at T4 and T5 with standardized math tests. Stronger N1 amplitudes for digits than false fonts were found across all 5 measurements. Arithmetic skills correlated negatively with visual N1 sensitivity to digits at T4 (2nd grade, mean age 8.3 yrs) over the left hemisphere, possibly reflecting allocation of more attentional or cognitive resources with poorer arithmetic skills. Our main result shows persistent visual N1 sensitivity to digits that is already present early on in pre-school and remains stable until fifth grade. This differs from the relatively sharp rise and fall of the visual N1 sensitivity to words or letters between kindergarten and middle of elementary school. The present study thus indicates different trajectories in the development of visual processing for written characters that are relevant to numeracy and literacy.

## 1 Introduction

Digits are symbols culturally associated with a representation of magnitude or quantity. Familiarization with numbers begins early in life, initializing basic processes that may have important consequences to the development of arithmetic and other academic skills (Bartelet *et al*., 2014; Lau *et al*., 2021). A frequently used model to describe number processing is the triple-code model (Dehaene, 1992; Dehaene *et al*., 2003), which proposes a set of separate systems for the representation of visual, verbal and quantity information. Each of these systems is supposed to rely on the specialization of sensory and assosiative brain areas (Dehaene, 2011), in a similar way as literacy leads to the development of brain networks for reading (Dehaene and Cohen, 2007). Understanding the development of specialized brain networks for digit and number processing could help characterize numeracy-related learning difficulties, such as dyscalculia (Butterworth, Varma and Laurillard, 2011).

A study using intracranial electrophysiological recordings identified a region in the inferior temporal gyrus showing preferential activation, i.e., stronger responses, to visual symbols denoting numerals compared to false fonts (Shum *et al*., 2013). The anatomical location of this *number form area* is close to another occipito-temporal area specialized for word processing in the fusiform gyrus (the *visual word form area*; Dehaene and Cohen, 2011; Yeatman, Rauschecker and Wandell, 2013). A recent study identified some character-specific regions in the left ventral occipito-temporal cortex using pseudowords and numbes (Rosenke *et al*., 2021). Of note, numerals in Shum et al., 2013 engaged different neuronal populations than words and letters. In sum, there is evidence supporting the notion that distinct cortical areas in the fusiform gyrus are specialized to process numerals and could explain why alexia patients with severe deficits in word and letter reading may have less impaired or even intact abilities to read numbers (see review in Starrfelt and Behrmann, 2011).

Interestingly, areas involved in both number and word processing in occipito-temporal cortex have been found to be preserved in congenitally blind people (Abboud *et al*., 2015). One possible explanation is that they may functionally speciliaze due to pre-existent structural connections with other regions associated with specific cognitive functions, rather than due to differences in the visual properties of the symbols and the underlying perceptual demands (Hannagan *et al*., 2015). In the case of areas specialized for numbers, there is evidence for functional connectivity with the intraparietal sulci (Abboud *et al*., 2015), associated with quantity representation and abstract magnitude processing (Piazza *et al*., 2004; Kadosh *et al*., 2011; Sasanguie *et al*., 2013). Similarly, the visual word form area is also functionally connected to core language areas (Stevens *et al*., 2017). Another non-exclusive hypothesis suggests that parts of occipito-temporal cortex may be sensitive to abstract properties of objects which can be encoded in different sensory modalities (e.g., soundscapes), which could explain the sensitivity observed also in blind subjects (Hannagan *et al*., 2015). This suggests that these areas belong to a hierarchically organized set of regions that are not exclusively responsible for visual processing. Thus, it would be important to consider additional properties (e.g., emerging arithmetic skills) when interpreting the development of visual responses to numbers to investigate top-down influences on number processing.

Number-specific subregions in the visual cortex have not been consistently detected by functional magnetic resonance imaging (fMRI) studies. Some meta-analyses suggested high convergence of functional activations in occipital areas (Arsalidou and Taylor, 2011; Sokolowski *et al*., 2017) but they did not differentiate between symbolic (e.g., Arabic) and nonsymbolic (e.g., dots) stimuli. Another meta-analysis took this into account and emphasized that differences in task demands across studies are crucial to understand the inconsistent findings (Yeo, Wilkey and Price, 2017). It has also been suggested that digits and letter strings may be represented as different patterns of activity in the visual cortex rather than recruiting separate regions (Peters, de Smedt and Op de Beeck, 2015). Despite these differences, neuroimaging evidence supports that regions in the posterior inferior temporal gyrus, part of ventral occipito-temporal cortex (vOTC), selectively respond stronger to numerals and number strings compared to other printed visual categories like words or formulas (e.g., Amalric and Dehaene, 2016; Grotheer, Herrmann and Kovács, 2016).

Previous cross-sectional evidence suggested a developmental shift in visual processing of letters and numbers from childhood (7- and 10-year olds) to adolescence (15-year-olds), characterized by an increase of visual responses to letters compared to digits (Park *et al*., 2018). However, it is yet to be investigated, how visual specialization to numbers develops in childhood, during the initial acquisition of numeric and arithmetic skills. A few fMRI studies have addressed the development of brain activation related to mathematical operations; they used explicit tasks involving calculations, number comparisons and thus probing more complex cognitive processing than the current study, which focuses on basic and implicit single digit processing: One of them examined symbolic and non-symbolic calculations and magnitude comparisons in children in third and sixth grades and in adults (Kucian *et al*., 2008). Although the networks engaged were similar, adults compared to children showed stronger activations in bilateral occipital and right inferior parietal regions during non-symbolic magnitude comparison (comparing the number of objects presented), and in the left inferior frontal gyrus during exact calculations. Children showed stronger activation in anterior cingulate gyrus relative to adults. A recent fMRI study compared activations in 10-12 year-old children and adults in several number comparison tasks (Skagenholt, Skagerlund and Träff, 2021). While a verbal comparison task yielded stronger activations of left-perysilvian language areas in adults, no group differences were found in the Arabic digit comparisons task. Focusing on children’s developmental data, another fMRI study examined brain activations longitudinally in 8-11 year-old children with and without developmental dyscalculia, at two time points during a period of 4 years (McCaskey *et al*., 2018). The study, which used a numerical order judgement task, found an increase in activation over time in frontal and parietal regions in children with dyscalculia, but a stable pattern of activations in controls. No significant differences in vOTC regions were found between the groups, but this could be due to limited statistical power and they were detected when lowering the rather strict significance threshold. In addition, stronger occipito-temporal activations in children with dyscalculia were previously reported in a study examining magnitude comparison of pictures (Kucian *et al*., 2011). These studies provide insights regarding brain networks for arithmetic, ordinal and quantity processing, but did not focus on the development of basic visual processing of numbers.

The current study uses electroencephalography (EEG) to investigate the development of visual sensitivity to single digits in children from pre-school and through elementary school, covering five time points. We focus on the visual N1, an electrophysiological correlate of vOTC activity. The N1 is an early component of the event-related potential (ERP) that peaks around 170 ms after stimuli presentation and is detected in posterior and bilateral occipito-temporal scalp electrodes. Besides general visual expertise (Tanaka and Curran, 2001), the amplitude of the N1 component has been associated with visual letter and word processing in multiple neuroimaging studies of reading and dyslexia (e.g., Maurer *et al*., 2006; Brem *et al*., 2013; Fraga González *et al*., 2014; Araújo *et al*., 2015). The EEG study from Park et al., (2018) examined N1 responses to strings of numbers, letters and false fonts in a cross-sectional design with 7-, 10-, 15-year-olds and young adults (Park *et al*., 2018). The results suggested a different trajectory emerging for a finer letter vs number contrast, compared to a coarser familiar vs unfamiliar contrast (i.e., letters or numbers vs false fonts). N1 amplitudes did not discriminate between letters and numbers in the younger 7-years-old group, but became larger to letters than to numbers in all older groups. Regarding the coarser discrimination, the two younger groups showed stronger responses for the familiar symbols but this pattern changed in the 15-years-old and adult groups, which showed stronger amplitudes for false fonts, in line with a previous report in adults from the same lab (Park *et al*., 2014). Park et al. (2018) provide an important reference to the development of visual responses to letters and numbers. However, their sparse time points do not cover the initial periods when letter and number knowledge, as well as arithmetic and reading skills emerge and rapidly change. The current study aims to clarify visual N1 specialization to numbers across five time points from preschool to fifth grade and focus on how arithmetic skills influences the development of the visual N1 to numbers.

In a recent publication, N1 sensitivity to single letters was compared to false fonts from pre-school to fifth grade (Fraga González *et al*., 2021)^1^. The results showed stronger N1 to letters than to false fonts in first grade, when children attain more advanced letter knowledge, but not in pre-school or in second or fifth grade. Thus, an inverted U-shaped development was suggested and interpreted within the predictive coding framework of vOTC specialization (Price and Devlin, 2011). This framework postulates that efficient visual processing is facilitated by top-down predictions, which emerge in early learning stages. This emergence is followed by strong prediction error signals indicating the mismatch between predicted and sensory inputs, which is used to optimize predictions (Friston, 2010). The combination between high prediction and prediction error signals at this stage of intensive learning explains the peak in neural responses. As learning progresses and expertise is attained, predictions become better and error signals decline, resulting in reduced activations. Alternatively, the inverted U trajectory can also be explained by principles of neural plasticity based on *expansion* (growth in number of neurons or connections involved), *selection* of most efficient circuits and subsequent *renormalization* when unnecessary circuits are pruned away (Wenger *et al*., 2017). These views on neural specialization and skill learning may also apply to learning numbers and arithmetic skills.

In the current study, larger N1 amplitudes to single digits compared to false fonts are considered to index vOTC sensitivity to digits. We aim to clarify the trajectory of this sensitivity across five time points from kindergarten through elementary school, covering the early and possibly most crucial period in learning basic number processing skills. To examine parallels with literacy acquisition, we include the analysis of N1 amplitudes to letters, which was the focus of previous work (Fraga-González et al., 2021). These developmental analyses are followed by an examination of the associations between N1 sensitivity to numbers and cognitive performance, with a special focus on arithmetic skills. The overall goal of this study is to understand how basic number processing is reflected in the specialization of visual areas and whether individual differences in visual processing of numerals are related to mathematical achievement.

## 2 Materials and Methods

### 2.1 Participants

The current study combines longitudinal and cross-sectional samples from a larger group of German-speaking children who took part in a large project with simultaneous EEG/fMRI sessions, behavioral tests and a grapheme-phoneme intervention training (Karipidis *et al*., 2017, 2018; Pleisch *et al*., 2019; Mehringer *et al*., 2020; Wang *et al*., 2020; Fraga González *et al*., 2021). Here, we focus on data from kindergarten (T1), middle of first grade (T2), end of first grade (T3), middle of second grade (T4), and middle of fifth grade (T5) of elementary school. Only data from participants meeting EEG data quality criteria in at least one time point for the experimental conditions digits and false fonts were analyzed (see section 2.5). In total, 62 participants fulfilled these criteria (35 female). Demographic information of this sample is presented in Table 1 and Table 2. An overview of the sample available at each time point is presented in Suppl. Fig. A.1. The number of participants available for each time point were 23, 22, 27, 27 and 42, for T1, T2, T3, T4 and T5, respectively. The number of participants with available data at one, two, three, four, and five time points were 27, 9, 10, 10 and 6, respectively. Familial risk for developmental dyslexia was estimated with the Adult Reading History Questionnaire (ARHQ; Lefly and Pennington, 2000). Individual risk scores were defined as the highest parental ARHQ value (Table 1). All participants had nonverbal IQ scores > 80, normal or corrected to normal visual acuity, and no neurological or cognitive impairments. Two participants had a diagnosis of ADHD (medication was interrupted 48h before test sessions), two participants reported having a sibling with reading impairments and one participant had adequate speech abilities during our study but a history of delayed speech development earlier in childhood. All children gave oral assent and parents gave written informed consent. The children received vouchers and presents as compensation for their participation. The project was approved by the local ethics committee of the Canton of Zurich and neighboring Cantons in Switzerland.

**Table 1.** Sample characteristics.

|  | *M (SD)[min,max]* |
| --- | --- |
| N | 62 |
| Sex ratio (male:female) | 27:35 |
| Handedness (right:left:both) | 56:6 |
| Nonverbal IQ <sup>a</sup> | 102.46 (8.80) [80, 121] |
| ARHQ<sup>b</sup> | 0.49 (0.16) [0.15, 0.80] |

<sup>a</sup>IQ was estimated from RIAS at T5 in 45 participants, from CFT at T4 in 16 children and from HAWIK in T1 for 1 participant. 5; <sup>b</sup>Familial risk level was low in 7 children (11.29 %, ARHQ < 0.3), moderate in 15 children (24.19 %, ARHQ range 0.3-0.4) and high in 40 children (64.51 %, ARHQ > 0.4).

**Table 2.** Descriptive statistics showing cognitive assessment scores at each test time.

|  | T1<br>(N = 23) | T2<br>(N = 22) | T3<br>(N = 27) | T4<br>(N = 27) | T5<br>(N = 42) |
| --- | --- | --- | --- | --- | --- |
|  | *M (SD)* | *M (SD)* | *M (SD)* | *M (SD)* | *M (SD)* |
| Age | 6.60 (0.52) | 7.38 (0.29) | 7.68 (0.30) | 8.28 (0.49) | 11.40 (0.41) |
| Sex (female : male) | 11:12 | 16:6 | 18:9 | 17:10 | 27:15 |
| Handedness (left : right) | 22:1 | 21:1 | 24:3 | 24:3 | 38:4 |
| Number-knowledge | 14.78 (3.78) | 18.36 (2.48) | 18.93 (2.40) | 20.74 (1.16) | - |
| HRT-Arithmetic skills raw |  |  |  |  |  |
| Addition | - | - | - | 16.15 (3.73) | 28.00 (5.15) |
| Subtraction | - | - | - | 14.78 (4.07) | 27.43 (5.64) |
| Completion | - | - | - | 8.19 (3.31) | 16.24 (5.88) |
| Comparison | - | - | - | 15.22 (4.52) | 29.23 (5.79) |
| Multiplication | - | - | - | 8.67 (3.50) | 24.17 (6.48) |
| Division | - | - | - | - | 24.22 (7.71) |
| Speeded writing | - | - | - | 19.63 (4.33) | 30.10 (5.45) |
| HRT-Arithmetic skills (PR) |  |  |  |  |  |
| Addition | - | - | - | 36.85 (24.47) | 39.05 ( 31.15) |
| Subtraction | - | - | - | 41.96 (24.89) | 46.91 ( 27.87) |
| Completion | - | - | - | 53.56 (26.68) | 48.57 ( 30.17) |
| Comparison | - | - | - | 55.15 (24.46) | 61.51 ( 27.73) |
| Multiplication | - | - | - | - | 35.66 ( 29.27) |
| Division | - | - | - | - | 42.76 ( 29.61) |
| Total Operations | - | - | - | 47.85 (27.10) | 46.18 ( 31.15) |
| Speeded writing | - | - | - | 46.15 (26.23) | 43.27 ( 25.87) |
| Letter-knowledge<sup>a</sup> |  |  |  |  |  |
| Sounds | 16.09 (10.48) | 45.55 (4.18) | 47.44 (2.94) | 49.37 (2.99) | - |
| SLRT-II reading fluency |  |  |  |  |  |
| Word |  | 7.09 (4.94) | 16.22 ( 9.64) | 25.85 (12.23) | 63.95 (24.13) |
| Pseudoword |  | 11.32 (6.42) | 17.44 (7.13) | 21.33 (7.31) | 39.98 (16.21) |
| Word (PR) | - | - | 55.37 (27.40) | 26.69 (22.30) | 31.82 (28.79) |
| Pseudoword (PR) | - | - | 50.22 (24.90) | 26.70 (19.57) | 34.07 (31.54) |
| Average (PR) | - | - | 52.80 (24.59) | 26.69 (20.14) | 32.95 (29.61) |

<sup>a</sup> Sum of scores for items in lower and upper case (max score = 52); PR = percentile score. Raw scores are: number of correct items (Number-knowledge, Letter-knowledge), number of correct items within 1 minute (SLRT-II).

### 2.2 Cognitive assessments

Cognitive tests were performed at each time point, depending on the test’s relevance to the specific grade level stage (Table 2). Number knowledge was assessed from T1 to T4 and the test consisted of naming twenty-one numbers, including all single digits from 1 to 9, and including numbers of up to three digits. Letter knowledge was assessed from T1 to T4 by asking participants to first identify the sound and then to name each letter from the (German) alphabet presented in blocks of upper and lower case letters (52 items in total). Basic arithmetic skills were assessed at T4 and T5 with several subtasks from the Heidelberger Rechentest (Haffner *et al*., 2005). The main subtasks were addition, subtraction, multiplication, comparison (fill gaps in simple formulas with the operators >, < or =), and completion (fill numbers in simple formulas, e.g., 6 = _ + 3). A subtask with divisions was also included in T5. There were no normative scores available for multiplication and division in T4. Each subtask consisted of 40 items and raw scores were computed based on the number of correct responses within 2 minutes, except for the comparison task in which half the number of errors was subtracted from the correct responses. A total operations percentile score was calculated from normative values of the individual operation subtests. In addition, the speeded writing subtask was used to assess visuomotor skills by requiring to copy numbers from a list of 60 items within 30 seconds.

Reading abilities were measured with the Salzburger Lese-und Rechtschreibtest (SLRT-II; Moll and Landerl, 2010). Fluency scores indicates the number of items correctly read within one-minute from lists of 156 items presented in eight columns. The lists could contain either words or pseudowords, and for each there were two lists available: List A was used in T1, T2 and T3, while list B was used in T4 and T5.

Finally, nonverbal IQ was estimated in T5 with the Reynolds Intellectual Assessment Scales (RIAS; Reynolds and Kamphaus, 2003), in T3 with the CFT1-R (Weiss and Osterland, 2013) and in T1 with the block design test of the Hamburg-Wechsler-Intelligenztest für Kinder (HAWIK-IV; Petermann and Petermann, 2010). The IQ estimate from the latest test time available was used (see note in Table 1). The description of rapid automated naming and phonological skills assessments are included in Appendix A, cognitive assessments.

### 2.3 Target detection task and stimuli

The current analysis focuses on an implicit audiovisual target detection task performed during simultaneous EEG-fMRI recordings. The complete paradigm was divided in four parts (at T1-T4) and three parts (at T5) of 375 s, each of them presenting unimodal visual and auditory, as well as bimodal stimulation blocks separated by fixation periods of 6 or 12 s. Each time point included the presentation of the following three character types in separate parts: letters, digits, and false fonts. Participants were instructed to press a button whenever a target stimulus (picture or sound of an animal/tool) appeared on the screen. The task is illustrated in Suppl. Fig. A.2. Our main analysis includes the unimodal visual conditions digits (DIG), letters (LET) and false fonts (FF), with a focus on the development of DIG vs FF differences. The analysis with a focus on LET vs. FF processing in a partially overlapping sample can be found in our previous publication (Fraga González *et al*., 2021). Each condition consisted of 4 blocks with 15 items per condition (in total 54 trials and 6 target trials per condition). In the DIG condition, the stimuli were the Arabic numerals 1-6 (all participants were able to name these numbers with 100% accuracy). In the LET condition, the stimuli included the letters *b, d, m, t, u, z* from the Latin alphabet presented in ‘Swiss school’ font (Fig. A.2.). In the FF condition, the stimuli were two sets of characters matched in size and width with the LET characters, created by rearranging different parts of those letters (Karipidis *et al*., 2017). All stimuli were visually presented using goggles (VisuaStimDigital, Resonance Technology, Northride, CA) in black in the middle of a grey background (mean visual angles horizontally/vertically DIG: 3°/6.7; FF: 2.8/4.8°). The stimuli in each block were presented pseudorandomized, with a duration of 613 ms and followed by an interestimulus interval of either 331 or 695ms. The task was programmed and presented using Presentation^®^ software (version 16.4, www.neurobs.com) and the design was adjusted to find a compromise between the optimal designs for EEG and fMRI recordings, and to account for the attentional demands of young children.

### 2.4 EEG data acquisition and preprocessing

This section largely overlaps with that of our recent report (Fraga-González et al., 2021). EEG data were recorded at 1 kHz sampling rate using an MR-compatible 128-channel EEG system (Net Amps 400, EGI HydroCelGeodesic Sensor Net) during functional magnetic resonance imaging (fMRI) in a Philips Achieva 3T scanner (Philips Medical Systems, Best, The Netherlands). Two additional electrodes placed on the chest registered the electrocardiogram (ECG). Impedances of the scalp electrodes were kept below 50 kΩ and the EEG system was synchronized with the scanner clock to reduce MR gradient artifacts. The recording reference was located at Cz and the ground electrode posterior to Cz. Vibrations were minimized by covering the electrodes with a bandage retainer net and turning off the helium pump and ventilation of the MRI scanner during image acquisition. Data were preprocessed with VisionAnalyzer 2.1 (BrainProducts GmbH, Munich, Germany). Electrodes with overall poor data quality or excessive artifacts were topographically interpolated (mean of 1.52 ±0.54 and no more than 5 electrodes in a subject). MR artifacts were removed using the average template subtraction method (Allen, Josephs and Turner, 2000) and ballistocardiogram artifacts were corrected using sliding average template subtraction as implemented in VisionAnalyzer 2.1. Additionally, continuous data were visually inspected to exclude periods with large artifacts like head movements. Subsequently, a band-pass filter of 0.1-30 Hz and 50 Hz Notch filter were applied and data were downsampled to 500 Hz. Then, we ran an independent component analysis (ICA) to remove components associated with blinks, eye movements, and residual ballistocardiogram artifacts. After artifact correction, data was visually inspected, electrodes located on the cheeks (E43, E48, E119, E120) were removed as they frequently contained major artifacts and a 0.1 Hz high-pass filter was applied to minimize residual slow artifacts. Finally, data were re-referenced to the common average reference.

### 2.5 Event-related potential analysis

The continuous EEG data were epoched from −100 to 613 ms after stimulus onset. Epochs with amplitudes ±200 μV or segments visually identified as containing residual artifacts were discarded from analysis. Only participants with at least 20 epochs in each condition were included in the analysis. The mean (SD) number of epochs for the DIG condition were T1: 40.91 (8.93), T2: 44.23 (9.41), T3: 42.96 (9.09), T4: 37.96 (9.94) and T5: 47.93 (5.58). For the FF condition they were T1: 43.87 (5.22), T2: 43.45 (8.56), T3: 46.48 (4.31), T4: 41.56 (10.69) and T5: 47.69 (5.10). There were no significant differences between conditions in the number of segments in any of the time points, *ps* > 0.077. The N1 time window was defined using the global field power (GFP; Lehmann and Skrandies, 1980) of the ERP averaged across both conditions and all subjects per time point. The interval was defined as ±30 ms around the GFP peak, i.e., around the second local maximum of the GFP, which corresponded with a typical N1 topography. The following intervals were defined independently at each time point to account for potential latency differences between measurements: T1 198-258 ms, T2: 188-248 ms, T3: 190-250 ms, T4: 180-240 ms and T5: 178-238 ms. Last, mean amplitudes of the N1 intervals were computed from the signal of the left occipitotemporal (LOT) cluster: E57, E58(=T5), E65, E70(=O1), E63, E64, E69, E68, E73 and the right occipitotemporal (ROT) cluster: E83(=O2), E90, E96(=T6), E100, E89, E95, E99, E88, E94. The clusters were defined based on visual inspection of topographies and previous studies (Pleisch *et al*., 2019).

### 2.6 Statistical analysis

The main analysis was performed by applying a linear mixed model (LMM) with a random intercept on N1 mean amplitudes and the fixed factors hemisphere (LOT, ROT), condition (DIG, LET, FF) and time point (T1, T2, T3, T4, T5). Separate LMM were subsequently also applied within each time point. The LMM was implemented with the function *lme* of the R package ‘Nlme’(Pinheiro *et al*., 2019). In our models, outliers were excluded if the normalized residuals exceeded the ± 3 threshold, which resulted in exclusion of 7 data points (1 % of the data) in the main model. Q-Q plots and predicted vs. residual plots were inspected to assess whether the data met the assumptions of normality and homoscedasticity. Differences in topography between conditions during the N1 intervals were examined at each time point with a topographical analysis of variance (TANOVA; Strik *et al*., 1998) implemented with in-house scripts and Matlab functions (R2017a, MathWorks, Natick, MA). At each time point, normalization of voltage amplitudes across condition were used to examine activity distribution and bootstrapping statistics were computed per data point (5000 permutations, as recommended for an accurate estimate of the significance at the 1% level; Manly, 2007). Finally, we used linear regressions to study the association between neural, i.e., N1 digit-false font amplitude differences at left and right hemisphere, and behavioral measures of interest, i.e., number knowledge at T1 to T3 and arithmetic and reading skills at T4 and T5 (SPSS Version 25.0. Armonk, NY: IBM Corp). The regressions were examined before and after exclusion of outliers in the neural measures of interest (a maximum of 4 cases were excluded in T3).

### 2.7 Data availability

Further information and requests for resources should be directed at and will be fulfilled by the Lead Contact, Silvia Brem. Some restrictions apply for data sharing for ethical reasons, because this would compromise participant confidentiality and privacy.

## 3 Results

### 3.1 Cognitive performance

The analysis of cognitive tests yielded the expected significant time effects suggesting improvements in the raw scores for the main assessments (see Fig. 1 and Table 2). Percentile scores indicating age-adjusted skill level are presented for the assessments and time points for which normative samples were available.

**Fig. 1.**
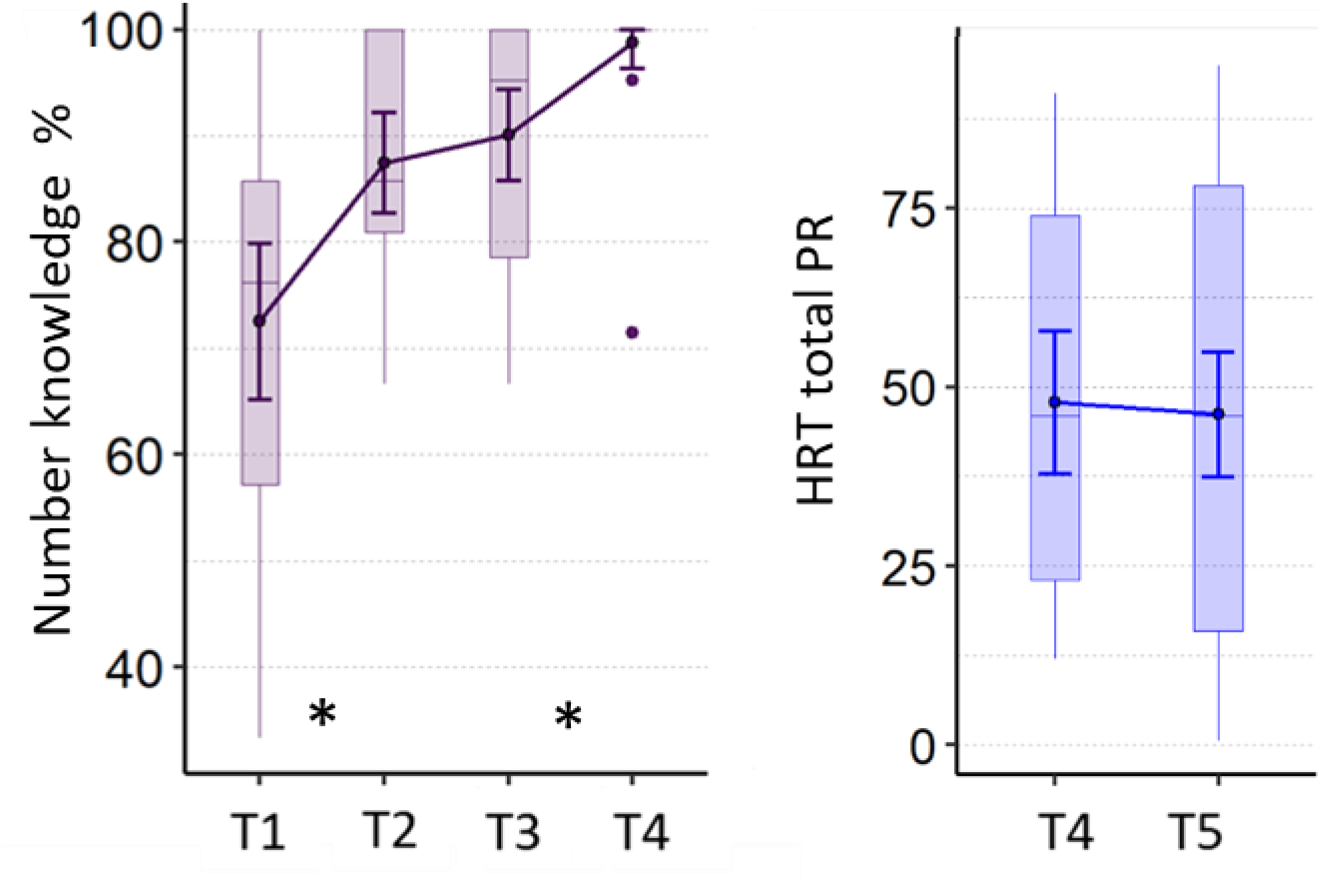
Numerical skills across measurements. Y-axis in left panel shows the percentage of correctly named numbers and in the right panel, the total percentile score for arithmetic operations. Asterisks indicate significant differences at *p* < .05 between test times. Boxplots show median and interquartile range. Error bars indicate mean and 95 % CIs.

Number knowledge increased significantly from T1 to T4, as revealed by the corresponding linear mixed model analysis on the effect of measurement time, *F* (3,53) = 45.13, *p* < 0.001. The scores reached ceiling levels at T4, and pairwise comparisons showed significant differences between T1-T2 (*t* (53) = −6.38, *p* < 0.001) and T3-T4 (*t* (53) = −3.77, *p* = 0.002), but not in the period of T2-T3, *p* = .652. A similar pattern was found for letter sounds knowledge, with main effect of time (*F* (3,53) = 350.8, *p* < 0.001) and significant gains in T1-T2 (*t* (53) = −23.92, *p* < 0.001), but not in the T2-T3 and T3-T4 periods, *ps* > 0.102. The individual arithmetic skills increased from T4 to T5, as revealed by a significant effect of time on the HRT tests of addition (*F* (1,17) =160.31, *p* < 0.001), substraction (*F* (1,17) = 151.57, *p* < 0.001), comparison (*F* (1,17) = 135.91, *p* < 0.001), completion (*F* (1,17) = 65.57, *p* < 0.001), and the additional speeded writing subtest (*F* (1,17) = 142.33, *p* < 0.001). There were no differences over time on the corresponding percentile scores, *ps* > 0.127. The number of participants performing < 16^th^ percentile in arithmetic operations (which may indicate poor math ability) in T1, T2, T3, T4 and T5 were 7, 5, 7, 8 and 12, respectively. Word and pseudoword reading were assessed from T2 to T5. Because different lists of items were used across time points (see section 2.2), we tested T2 vs T3 and T4 vs T5 separately. The analysis showed a significant increase between T2 and T3 in the number of correctly read words (*F* (1,16) = 26.35 *p* < 0.001) and pseudowords (*F* (1,16) = 22.23, *p* < 0.001). The same pattern was found between T4 and T5 for words (*F* (1,17) = 119.18, *p* < 0.001) and pseudowords (*F* (1,17) = 68.52, *p* < 0.001). The percentile scores showed no statistically significant differences between the test times, *ps* > 0.107. Finally, there were also significant gains in RAN and phonological skills. These analyses are reported in Appendix B (section 9.1) supplementary results, as they are not the focus of the current study.

### 3.2 ERP analysis

Individual condition-wise visual N1 mean amplitude to digits, letters and false fonts in the target detection task for each time point was used for statistical analyses. The current analysis focuses on the differences between digits and false fonts and also includes the comparison with letters (for an analysis focused on letter vs false fonts in a partially overlapping sample see Fraga-González et al., 2021).

#### 3.2.1 Development of N1 responses to digits

##### 3.2.1.1 N1 mean amplitude

The ERP, topographies and GFP waveforms per time point (T1 to T5), condition (DIG, LET, FF) and electrode cluster (LOT, ROT) are shown in Fig. 2 (panels a, b and c). We performed a LMM analysis on N1 mean amplitudes with the factors time, condition and hemisphere (see sections 2.5 and 2.6 for N1 time range selection and model details). The analysis revealed a main effect of time point (*F* (4,748) = 8.18, *p* < 0.001), indicating significant differences between the time points across conditions and hemispheres. Moreover, there was a main effect of condition (*F* (1,746) = 63.72, *p* < 0.001), indicating differences in amplitudes between the conditions over all time points and hemispheres. In addition, there was a significant interaction between the factors time point and condition (*F* (8,746) = 2.40, *p* = 0.015), suggesting that these differences changed over time. No other effects were statistically significant, but there was a trend for hemisphere effects, at *p* = 0.090. The mean amplitudes per condition, time point and hemisphere are shown in Fig.3. A more detailed figure with box plots, individual data points and distribution is presented in the supplementary figure B.1. This analysis was followed by LMMs for each pair of conditions.

**Fig. 2.**
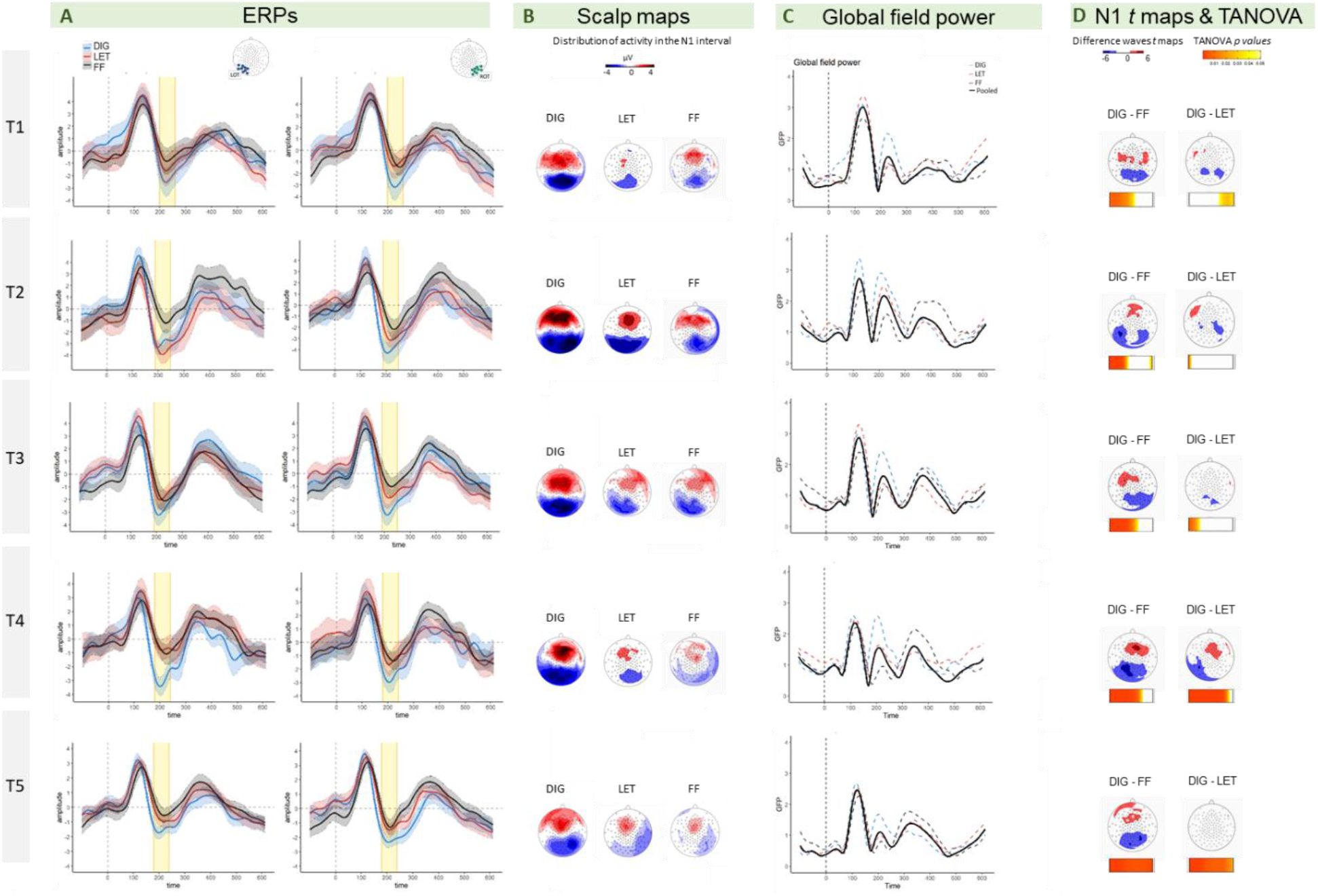
*Panel A*. ERPs (μV) for digits (blue lines), letters (red lines) and false fonts (black lines) with ribbons indicating 95% CIs (within-subject). Left hemispheric amplitudes are depicted on the left side and right hemispheric amplitudes on the right. The N1 time-window is highlighted in yellow. *Panel B*. Scalp topographies for digits, letters and false fonts. *Panel C*. GFP for the average of the three conditions (black line), digits (dashed blue line), letters ( dashed red line) false fonts (dashed black line). *Panel D*. *T*-test maps for the difference between conditions and TANOVA *p* values from permutation tests of distribution differences across the N1 time interval. ERPs = event-related potentials; CIs = confidence intervals; GFP = global field power; TANOVA = topographic analysis of variance; DIG = digits; LET = letters; FF = false fonts; LOT= left occipito-temporal; ROT = right occipito-temporal.

**Fig. 3.**
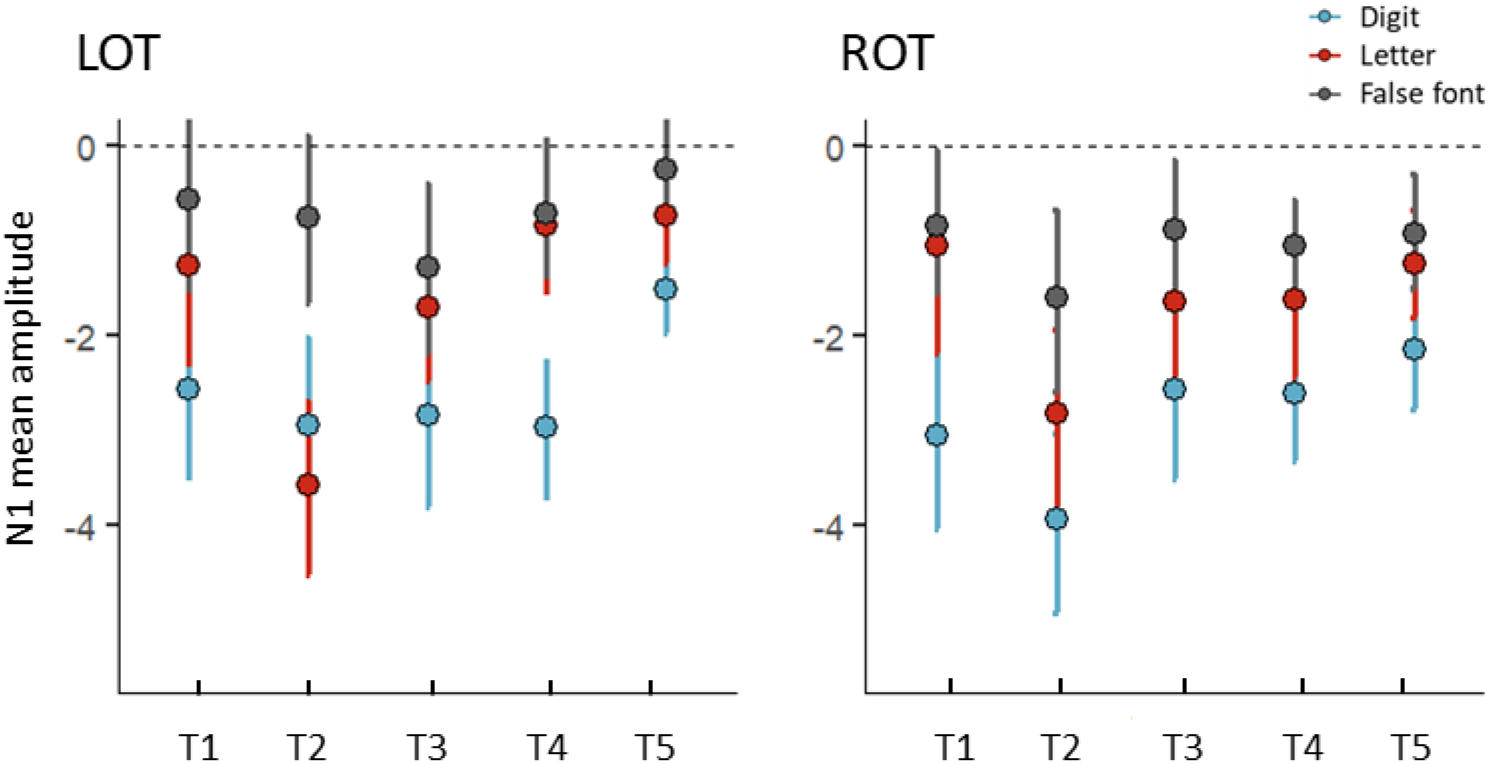
Mean amplitudes for the N1 interval over time points (T1,T2,T3,T4,T5) for digits (blue), letters (red) and false fonts (black). Left and right clusters in separate panels. Error bars represent 95 % CI. Normalized residuals from the main model exceeding ± 3 threshold were excluded (7 data points).

The LMM using digits and false fonts showed a effect of time point (*F* (4,480) = 3.22, *p* = 0.013), and a significant effect of condition (*F* (4,480) = 113.80, *p* < 0.001), but no interactions, *ps* > 0.111. Follow up tests per time point confirmed stronger amplitudes for digits vs false fonts in all time points (T1, *t* = −4.79, *p* < 0.001; T2, *t* = −5.40, *p* < 0.001; T3, *t* = −4.41, *p* < 0.001; T4, *t* = −5.05, *p* < 0.001; T5, *t* = −4.15, *p* < 0.001). The additional LMM with digits and letters showed main effects of time (*F* (4,476) = 10.08, *p* < 0.001) and condition (*F* (4,476) = 41.36, *p* < 0.001), and a trend for an interaction between time and condition, (*F* (4,476) = 2.13, *p* = 0.076). The contrasts of digits vs letters revealed significantly stronger amplitudes for digits in T1 (*t* = −4.00, *p* < 0.001), T3 (*t* = −2.58, *p* = 0.010), T4 (*t* = −4.35, *p* < 0.001) and T5 (*t* = −3.08, *p* = 0.002), but not in T2 (*p* = 0.545). The LMM with letters and false fonts (see also Fraga-González et al., 2021), showed main effects of time (*F* (4,477) = 6.22, *p* < 0.001) and condition (*F* (4,477) = 23.03, *p* < 0.001), together with an interaction between time and condition (*F* (4,477) = 3.73, *p* = 0.005). The follow-up tests per time point comparing letters vs false fonts only revealed stronger amplitudes for letters in T2, *t* = −5.31, *p* < 0.001, all other *ps* < .151. Additional topographical maps of t-tests on the difference between each pair of conditions are presented in Fig. 2 (panel d). The figure shows a mainly bilateral distribution of the digit minus false font difference. Exploratory post-hoc comparisons per hemisphere and test time showed that this digit-false font difference was significant across time and hemisphere. The *t*-maps of the digit-letter differences showed a less defined topography, and suggest a potential left lateralization at T4. Exploratory follow-up comparisons supported this by showing a significant digit-letter difference at T4 in the left hemisphere (*t* = −4.10, *p* < 0.001) but not in the right hemisphere (*p* = 0.150). This result should, however, be interpreted with caution since the main model within the electrode cluster of interest did not reveale significant interactions with the factor hemisphere.

Further details of the N1 differences between digits and false fonts, which is focus of the current study, are presented in Fig.4. The percentage of participants showing stronger negativity for digits compared to false fonts in the left hemisphere were (T1) 82.61%, (T2) 77.27 %, (T3) 62.96%, (T4) 77.78 % and (T5) 73.81%. In the right hemisphere the percentages were (T1) 86.96 %, (T2) 77.27 %, (T3) 85.19 %, (T4) 74.07 % and (T5) 78.57 %. Regarding the digits minus letter difference, the percentage of participants with stronger N1 amplitudes for digits were (T1) 60.87 %, (T2) 31.82 %, (T3) 66.67 %, (T4) 84.62 % and (T5) 76.19 %; in the right hemisphere the percentages were (T1) 78.26 %, (T2) 63.64 %, (T3) 74.07 %, (T4) 69.23 % and (T5) 71.43 %. The distributions of the digits minus letter amplitudes per time point and hemisphere are presented supplementary figure B.2.

**Fig. 4.**
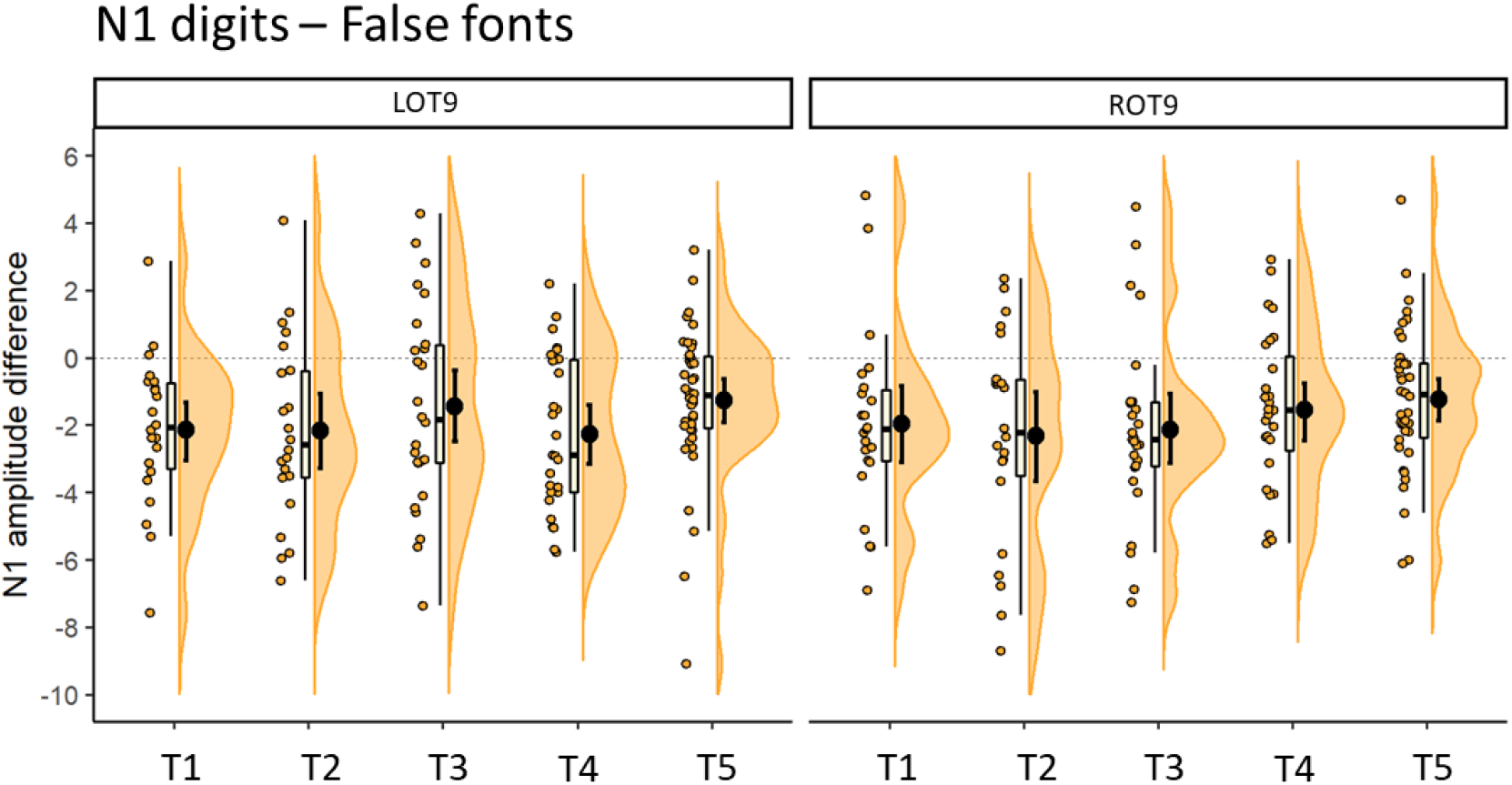
Differences in N1 amplitudes between digits and false fonts per time point. Error bars within the density plot indicate mean and 95 % CIs. More negative values indicate stronger negative amplitudes for digits than false fonts. Left hemispheric cluster is depicted on the left panel, right hemispheric cluster on the right.

##### 3.2.1.2 Topographic analysis of variance (TANOVA)

An exploratory TANOVA compared the topographical distribution of activity between conditions at each time point. Amplitudes were normalized in this analysis so the comparisons indicate differences in distribution independent from intensity of activity. The TANOVA results for the N1 time interval are plotted in Fig. 2 panel D. Significant differences in topography are indicated by *p* values from the permutation tests < 0.05. The comparison of digits and false fonts showed broad intervals of statistical significance around and after the N1 time window, and some less consistent periods preceding stimulus onset in T1 and T5. Regarding digits and letters, these periods are much more narrow centered on the N1 interval. Note that normalized activation maps were used in this analysis, thus these differences can be interpreted as independent from intensity of activity.

#### 3.2.2 N1 sensitivity to digits and cognitive performance

The following analyses were performed separately with left and right hemisphere amplitudes. We first examined whether digit sensitivity reflected by N1 digit-false font differences in kindergarten (T1) could predict later arithmetic or reading skills. Specifically, we focused on the latest score available (T5 when available, T4 otherwise) for the summary measure of arithmetic operations (HRT) and the mean of word and pseudoword reading (SLRT-II). The analyses revealed no significant associations between N1 digit sensitivity in T1 and later arithmetic or reading skills, *ps* > 0.144.

Next, the linear relationship between N1 digit sensitivity and arithmetic skills was investigated within the different learning stages. The core cognitive measures (predicted variables) were number knowledge in the first three time points (T1, T2, and T3) and arithmetic operations in 2^nd^ and 5^th^ grade (T4 and T5). The analysis on number knowledge yielded no statistically significant evidence for an association with N1 digit sensitivity in any of the three time points from T1 to T3, *ps* > 0.177. Here, it should be noted that performance in the number knowledge task was relatively high already in kindergarten and close to ceiling in subsequent measurements (see Fig.1a). The analysis on arithmetic skills yielded significant associations between the age-adjusted level of arithmetic skills and N1 in T4 (2^nd^ grade; N = 27). Specifically, the digit sensitivity in N1 amplitudes in the left hemisphere showed a significant relation with the HRT percentile summary score for arithmetic operations (*R* = 0.60, *R^2^* = 0.36, β = 7.02, *t* = 3.72, *p* = 0.001). This result is presented in Fig. 5. The direction of this relationship indicates larger digit sensitivity over the left electrode cluster in children with lower performance. In addition, this result was also statistically significant for the percentile scores of addition, (*R* = 0.60, *R^2^* = 0.36, β = 0.60, *t* = 3.74, *p* < 0.001), subtraction (*R* = 0.47, *R^2^* = 0.22, β = 0.47, *t* = 2.66, *p* = 0.013), comparison (*R* = 0.44, *R^2^* = 0.19, β = 0.44, *t* = 2.46, *p* = 0.021) and completion (*R* = 0.56, *R^2^* = 0.31, β = 0.55, *t* = 3.34, *p* = 0.003). There was no evidence for statistically significant associations with number writing speed, *p* = 0.285. Interestingly, no significant associations were found between HRT arithmetic operations and N1 digit-false font difference in the right hemisphere, *ps* > 0.269. The analysis in T6 yielded no significant associations, *ps* > 0.096. The same pattern of results was found when using the raw scores. This analysis is presented in Appendix B section 9.4.

**Fig.5.**
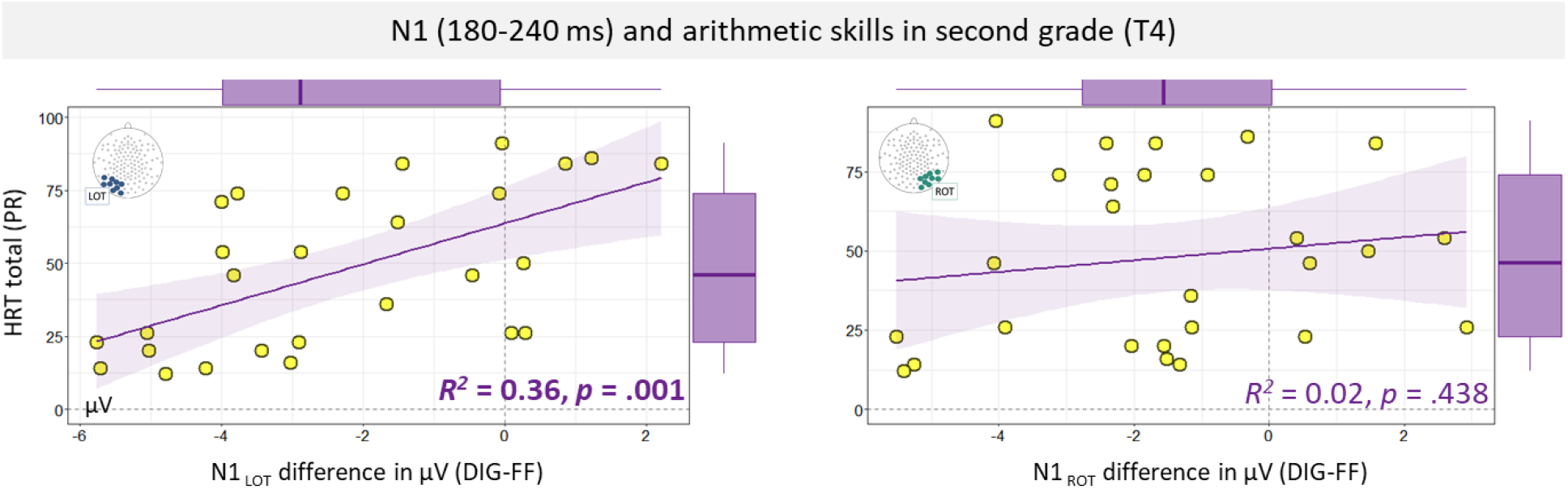
Linear regression with total percentile score of arithmetic operations (HRT total) as dependent variable and N1 digits-false font amplitude differences in the left and right ventral-occipital electrodes as predictor. Box plots are displayed in the margins. LOT = left occipio-temporal; ROT = right occipito-temporal; DIG = digits; FF = false fonts; HRT = Heidelberger Rechentest; PR = percentile.

## 4 Discussion

The current study focused on N1 amplitude differences between digits and false fonts across 5 time points in pre-school, middle and end of first grade, second grade and fifth grade. Additionally, associations between N1 digit sensitivity (indicated by larger N1 amplitudes to digits vs false fonts) and arithmetic and reading skills were also investigated. The results suggest N1 digit sensitivity that was present in all time points and correlated with arithmetic skills in 2^nd^ grade.

The main analysis revealed stronger N1 amplitudes to digits than false fonts across all time points from pre-school to fifth grade. The strength of this sensitivity seemed stable across time, as suggested by the ERPs, and the follow up analyses showing significant digit vs false font differences at each time point. This result contrasts with the development of N1 sensitivity to letters, which was examined in detail in a previous report (Fraga González *et al*., 2021). Letter vs false font sensitivity showed an inverted U-shaped trajectory; N1 showed sensitivity to letters when children attained letter knowledge in first grade (T2) and shortly after (T3), but not in pre-school or in second and fifth grade. The current analysis shows that digits elicited stronger amplitudes than letters in all time points except at T2.

Although digits and letters may be comparable in terms of visual features and levels of exposure, they diverge in several attributes which may critically contribute to the development of brain systems for numeracy and literacy. Digits start being learned earlier than letters and convey a wide range of information, some of which may only be assimilated over long years of learning, e.g., complex mathematical properties. In the earlier years of numeracy, such as those covered by the current study, aspects like numerosity and parity become increasingly important over the years (Miller and Gelman, 1983; Moeller *et al*., 2011). Importantly, these properties vary in the extent they may be automatically activated by different tasks (e.g., Cohen, 2009; Kallai and Tzelgov, 2012). Therefore, digits in the current paradigm could be co-activating representations of different semantic properties and these may differ over the time points, for example, when arithmetic skills emerge. According to the triple-code model of number processing, the non-verbal semantic representations of numbers, that is, the quantity system, would mostly recruit parietal areas, in particular the intraparietal sulcus region (Dehaene *et al*., 2003). In this model, a verbal system processing phonological and lexical representations of numbers would activate language areas, like the left angular gyrus. Recruitment of these areas may be shared between letters and digits, as well as domain-general areas (e.g., visuo-spatial attention) required to perform a specific task. However, we would expect a network associated with the quantity system to be specific to digits. A recent meta-analysis reviewed the triple-code model and suggested the inclusion of additional regions that are likely to be detected in calculation or number tasks, such as the cingulate gyri, insula and cerebellum (Arsalidou and Taylor, 2011). This metaanalysis also suggested a contribution of prefrontal areas involved in working memory processes and task difficulty.

If we consider the development of vOTC activations from an interactive perspective (Price and Devlin, 2011), it is possible that the interplay between specific number processing systems and domain general systems contribute to the current trajectory of digits, that diverges from the inverted U-shaped trajectory for letters. Previous studies showed that N1 can be affected by general task demands and focus of attention (Luck, Woodman and Vogel, 2000; Vogel and Luck, 2000; Yoncheva *et al*., 2010; Okumura, Kasai and Murohashi, 2015). More specifically, paying attention to nonspatial features (e.g., shape, color) can modulate visual ERPs in time windows such as that of N1 or earlier (Hillyard and Anllo-Vento, 1998; Yoncheva *et al*., 2010). In future studies, manipulation over the different number properties elicited by the experimental task could help clarifying these contributions to the N1 component.

The current digit vs false font data does not fit an inverted U-shaped trajectory that can be accommodated in the predictive coding framework for learning (Price and Devlin, 2011) or the cortical expansion-selection-renormalization model of skill learning (Wenger *et al*., 2017). An important point to discuss in relation to this, is the time period examined in the current study and the numeracy learning milestones captured by it. The results show that N1 sensitivity to digits can be detected already at T1, at kindergarten age. As shown in Fig 1, knowledge of numbers is already high at this stage, even though there is a substantial improvement from T1 to T2. Consequently, the current data do not allow us to assess the emergence of N1 digit sensitivity with initial attainment of number knowledge, which presumably happens earlier than the current T1 measurement. The single-digits in our paradigm were all well known to the participants already at T1. The lack of significant associations between N1 amplitudes and number knowledge may thus be due to the advanced knowledge of digits at T1. Interestingly, our results do not suggest a decay in this sensitivity in the subsequent time points, contrasting previous findings for words and letters (Maurer *et al*., 2006; Fraga González *et al*., 2021). Here, we should note that arithmetic operations skills are still intensively trained at T4 and T5. This period coincides with the outset of more fluent word and pseudoword decoding (see Table 2). It is possible that visual processing of digits remains functionally important to perform these operations and that there is also automatic processing of new properties which are learned in parallel with arithmetic skills (e.g., numerosity, parity). Single letters on the other hand, may elicit weaker visual responses as at this stage they are mostly utilized as part of larger units like syllables and words, which carry much more complex phonological and semantic information. The difference between a single digit and a multi-digit number is not as large as between single letters and letter strings in terms of their associations. This could explain the sustained N1 sensitivity to digits in this period, in contrast with the observed decline of N1 letter sensitivity (Fraga González *et al*., 2021).

We found a significant association between performance of simple arithmetic operations and N1 digit sensitivity at T4, but not at T5. Children with poorer arithmetic skills showed stronger N1 amplitudes for digits than false fonts over the left posterior electrodes. This may indicate that other systems for number processing, rather than visual, may be gaining functional importance as arithmetic skills and knowledge become more complex, i.e., the quantity or verbal system in the triple-code model (Dehaene, 1992). Later time points with corresponding measures of math performance would be useful to further investigate this. In addition, a visual N1 study on children, adolescents and young adults showed differences in 15-year-olds and adults compared to 7-and 10-year-olds in a coarse N1 sensitivity to letter and numbers compared to unfamiliar false fonts (Park *et al*., 2018). Park et al. (2018) used strings instead of single characters which makes the direct comparison with our analysis difficult. Nonetheless, the results of Park et al. support the relevance of examining later periods in development and also raise the question of the role of visual familiarity. A challenge here would be to examine N1 digit sensitivity by using control stimuli that can be matched in terms of visual familiarity (e.g., comparable level of exposure before performing the experimental task). Additionally, we cannot rule out that developmental changes in the attentional factors discussed above may also have contributed to these results.

Another point for discussion is N1 lateralization. The relation between N1 and arithmetic operations at T4 was only found in the left hemispheric cluster, which suggests functional interhemispheric differences in N1 digit sensitivity. This could reflect some interaction between left vOTC and left lateralized language or parietal areas that are instrumental to arithmetic operations. But it should be noted that the ERP topographies and lack of lateralization effects from the main model suggest that N1 digit sensitivity itself was bilateral. The previous developmental report on N1 to number strings suggested a late shift in lateralization towards more right lateralized N1 amplitudes for numbers that emerged in 15-year-olds and young adults (Park *et al*., 2018), which the current data did not capture. However, from previous findings on laterality of N1 to single characters it still remains unclear when this lateralization develops (Tarkiainen *et al*., 1999; Wong *et al*., 2005; Stevens *et al*., 2013; Fraga González *et al*., 2021).

A potential limitation regarding generalizability of the current findings is that our sample consists of children with varying degree of reading skills and familial risk for dyslexia. There is evidence for comorbidity between dyslexia and dyscalculia and for shared features in the development of language and math skills (Butterworth, Varma and Laurillard, 2011; Peng *et al*., 2020). Nonetheless, this also presents the advantage of capturing a wider range of math achievement as it was found in our sample. Finally, as discussed above, the current visual target detection task cannot account for differences in attentional strategies and automatic processing of semantic features during performance.

### 4.1 General conclusion

We examined the development of visual specialization for digits, reflected by stronger N1 ERP responses to digits than false fonts. The results show that visual N1 sensitivity to digits is present already in pre-school and remains stable through primary school. Visual processing of digits seems to be important at the beginning of arithmetic skill acquisition as suggested by the regression results showing an association between N1 amplitudes and math skills. Further research should clarify when this basic visual sensitivity to digits emerges and how it interacts with verbal and quantity systems for numerical cognition.

## 5 Acknowledgments

We thank C. Hofstetter, M. Röthlisberger, A. Brem, C. Brauchli, L. Götze, R. Rossi, F. Aepli, V. Keller, C. Wick, R. Füzér, M. Schneebeli, D. Dornbierer, M. Hartmann, S. Suter, T. Aegeter, L. Barblan, F. Mergen-Felten, L. Meyer, L. Vogel and P. Stämpfli for their assistance during recruitment, investigation, data acquisition and data analyses. We also thank all the participating children with their families.

## Footnotes

1 The sample of Fraga-González 2021 partially overlaps that of the current study.

## References

Abboud, S. et al. (2015) ‘A number-form area in the blind’, Nature Communications. Nature Publishing Group, 6(1), pp. 1–9. doi: 10.1038/ncomms7026.

Allen, P. J., Josephs, O. and Turner, R. (2000) ‘A method for removing imaging artifact from continuous EEG recorded during functional MRI’, NeuroImage. Academic Press Inc., 12(2), pp. 230–239. doi: 10.1006/nimg.2000.0599.

Amalric, M. and Dehaene, S. (2016) ‘Origins of the brain networks for advanced mathematics in expert mathematicians’, Proceedings of the National Academy of Sciences of the United States of America. National Academy of Sciences, 113(18), pp. 4909–4917. doi: 10.1073/pnas.1603205113.

Araújo, S. et al. (2015) ‘Lexical and sublexical orthographic processing: An ERP study with skilled and dyslexic adult readers’, Brain and Language, 141, pp. 16–27. doi: 10.1016/j.bandl.2014.11.007.

Arsalidou, M. and Taylor, M. J. (2011) ‘Is 2+2=4? Meta-analyses of brain areas needed for numbers and calculations’, NeuroImage, 54(3), pp. 2382–2393. doi: 10.1016/j.neuroimage.2010.10.009.

Bartelet, D. et al. (2014) ‘What basic number processing measures in kindergarten explain unique variability in first-grade arithmetic proficiency?’, Journal of Experimental Child Psychology. Academic Press, 117(1), pp. 12–28. doi: 10.1016/j.jecp.2013.08.010.

Brem, S. et al. (2013) ‘An electrophysiological study of print processing in kindergarten: the contribution of the visual N1 as a predictor of reading outcome.’, Developmental neuropsychology, 38(8), pp. 567–94. doi: 10.1080/87565641.2013.828729.

Butterworth, B., Varma, S. and Laurillard, D. (2011) ‘Dyscalculia: From brain to education’, Science. Science, pp. 1049–1053. doi: 10.1126/science.1201536.

Cohen, D. J. (2009) ‘Integers do not automatically activate their quantity representation’, Psychonomic Bulletin and Review. Springer, 16(2), pp. 332–336. doi: 10.3758/PBR.16.2.332.

Dehaene, S. (1992) ‘Varieties of numerical abilities’, Cognition. Elsevier, 44(1–2), pp. 1–42. doi: 10.1016/0010-0277(92)90049-N.

Dehaene, S. et al. (2003) ‘Three parietal circuits for number processing’, Cognitive Neuropsychology. Taylor & Francis Group, pp. 487–506. doi: 10.1080/02643290244000239.

Dehaene, S. (2011) The Number Sense: How the Mind Creates Mathematics, Revised and Updated Edition. Edited by OUP USA.

Dehaene, S. and Cohen, L. (2007) ‘Cultural recycling of cortical maps.’, Neuron, 56(2), pp. 384–98. doi: 10.1016/j.neuron.2007.10.004.

Dehaene, S. and Cohen, L. (2011) ‘The unique role of the visual word form area in reading.’, Trends in cognitive sciences. Elsevier Ltd, 15(6), pp. 254–62. doi: 10.1016/j.tics.2011.04.003.

Fraga González, G. et al. (2014) ‘Brain-potential analysis of visual word recognition in dyslexics and typically reading children.’, Frontiers in human neuroscience, 8(June), p. 474. doi: 10.3389/fnhum.2014.00474.

Fraga González, G. et al. (2021) ‘The rise and fall of rapid occipito-temporal sensitivity to letters: Transient specialization through elementary school’, Developmental Cognitive Neuroscience. Elsevier, 49, p. 100958. doi: 10.1016/j.dcn.2021.100958.

Friston, K. (2010) ‘The free-energy principle: A unified brain theory?’, Nature Reviews Neuroscience, pp. 127–138. doi: 10.1038/nrn2787.

Grotheer, M., Herrmann, K. H. and Kovács, G. (2016) ‘Neuroimaging evidence of a bilateral representation for visually presented numbers’, Journal of Neuroscience. Society for Neuroscience, 36(1), pp. 88–97. doi: 10.1523/JNEUROSCI.2129-15.2016.

Haffner, J. et al. (2005) Heidelberger Rechentest (HRT 1-4). Erfassung mathematischer Basiskompetenzen im Grundschulalter. Hogrefe, Hogrefe Schultests. Hogrefe.

Hannagan, T. et al. (2015) ‘Origins of the specialization for letters and numbers in ventral occipitotemporal cortex’, Trends in Cognitive Sciences. Elsevier Ltd, pp. 374–382. doi: 10.1016/j.tics.2015.05.006.

Hillyard, S. A. and Anllo-Vento, L. (1998) ‘Event-related brain potentials in the study of visual selective attention’, Proceedings of the National Academy of Sciences. National Academy of Sciences, 95(3), pp. 781–787. doi: 10.1073/PNAS.95.3.781.

Kadosh, R. C. et al. (2011) ‘Specialization in the human brain: The case of numbers’, Frontiers in Human Neuroscience. Frontiers Media S. A., 5(JULY), p. 62. doi: 10.3389/fnhum.2011.00062.

Kallai, A. Y. and Tzelgov, J. (2012) ‘The place-value of a digit in multi-digit numbers is processed automatically’, Journal of Experimental Psychology: Learning Memory and Cognition, 38(5), pp. 1221–1233. doi: 10.1037/a0027635.

Karipidis, I. I. et al. (2017) ‘Neural initialization of audiovisual integration in prereaders at varying risk for developmental dyslexia’, Human Brain Mapping, 38(2), pp. 1038–1055. doi: 10.1002/hbm.23437.

Karipidis, I. I. et al. (2018) ‘Simulating reading acquisition: The link between reading outcome and multimodal brain signatures of letter–speech sound learning in prereaders’, Scientific Reports. Nature Publishing Group, 8(1), p. 7121. doi: 10.1038/s41598-018-24909-8.

Kucian, K. et al. (2008) ‘Development of neural networks for exact and approximate calculation: A fMRI study’, Developmental Neuropsychology. Taylor & Francis Group, 33(4), pp. 447–473. doi: 10.1080/87565640802101474.

Kucian, K. et al. (2011) ‘Non-symbolic numerical distance effect in children with and without developmental dyscalculia: A parametric fMRI study’, Developmental Neuropsychology. Taylor & Francis Group, 36(6), pp. 741–762. doi: 10.1080/87565641.2010.549867.

Lau, N. T. T. et al. (2021) ‘Kindergarteners’ symbolic number abilities predict nonsymbolic number abilities and math achievement in grade 1.’, Developmental Psychology. American Psychological Association (APA), 57(4), pp. 471–488. doi: 10.1037/DEV0001158.

Lefly, D. L. and Pennington, B. F. (2000) ‘Reliability and Validity of the Adult Reading History Questionnaire’, Journal of Learning Disabilities, 33(3), pp. 286–296. doi: 10.1177/002221940003300306.

Lehmann, D. and Skrandies, W. (1980) ‘Reference-free identification of components of checkerboard-evoked multichannel potential fields’, Electroencephalography and Clinical Neurophysiology, 48(6), pp. 609–621. doi: 10.1016/0013-4694(80)90419-8.

Luck, S. J., Woodman, G. F. and Vogel, E. K. (2000) ‘Event-related potential studies of attention.’, Trends in cognitive sciences, 4(11), pp. 432–440. Available at: http://www.ncbi.nlm.nih.gov/pubmed/11058821.

Manly, B. F. J. (2007) Randomization, bootstrap and Monte Carlo methods in biology. Boca Raton, FL: Chapman & Hall/CRC.

Maurer, U. et al. (2006) ‘Coarse neural tuning for print peaks when children learn to read.’, NeuroImage, 33(2), pp. 749–58. doi: 10.1016/j.neuroimage.2006.06.025.

Mayer, A. (2011) ‘Test zur Erfassung der phonologischen Bewusstheit und der Benenngeschwindigkeit (TEPHOBE)’. Ernst Reinhardt Verlag, München.

McCaskey, U. et al. (2018) ‘Longitudinal brain development of numerical skills in typically developing children and children with developmental dyscalculia’, Frontiers in Human Neuroscience. Frontiers Media S. A, 11, p. 629. doi: 10.3389/fnhum.2017.00629.

Mehringer, H. et al. (2020) ‘(Swiss) GraphoLearn: an app-based tool to support beginning readers’, Research and Practice in Technology Enhanced Learning, 15(1), pp. 1–21.

Miller, K. and Gelman, R. (1983) ‘The Child’s Representation of Number: A Multidimensional Scaling Analysis’, Child Development. JSTOR, 54(6), p. 1470. doi: 10.2307/1129809.

Moeller, K. et al. (2011) ‘Early place-value understanding as a precursor for later arithmetic performance-A longitudinal study on numerical development’, Research in Developmental Disabilities. Pergamon, 32(5), pp. 1837–1851. doi: 10.1016/j.ridd.2011.03.012.

Moll, K. and Landerl, K. (2010) ‘SLRT-II: Lese-und Rechtschreibtest’. Huber, Bern.

Okumura, Y., Kasai, T. and Murohashi, H. (2015) ‘Attention that covers letters is necessary for the left-lateralization of an early print-tuned ERP in Japanese hiragana’, Neuropsychologia. Elsevier, 69, pp. 22–30. doi: 10.1016/j.neuropsychologia.2015.01.026.

Park, J. et al. (2014) ‘Experience-dependent hemispheric specialization of letters and numbers is revealed in early visual processing’, Journal of cognitive neuroscience, 26(10), pp. 2239–2249. doi: 10.1162/jocn_a_00621.

Park, J. et al. (2018) ‘Developmental trajectory of neural specialization for letter and number visual processing’, Developmental Science. Blackwell Publishing Ltd, 21(3), p. e12578. doi: 10.1111/desc.12578.

Peng, P. et al. (2020) ‘Examining the mutual relations between language and mathematics: A meta-analysis’, Psychological Bulletin. American Psychological Association Inc., 146(7), pp. 595–634. doi: 10.1037/bul0000231.

Petermann, F. and Petermann, U. (2010) ‘HAWIK-IV: Hamburg-Wechsler-Intelligenztest für Kinder-IV; Manual; Übersetzung und Adaption der WISC-IV von David Wechsler’. Huber.

Peters, L., de Smedt, B. and Op de Beeck, H. P. (2015) ‘The neural representation of arabic digits in visual cortex’, Frontiers in Human Neuroscience. Frontiers Media S. A, 9(September), p. 517. doi: 10.3389/fnhum.2015.00517.

Piazza, M. et al. (2004) ‘Tuning curves for approximate numerosity in the human intraparietal sulcus’, Neuron. Cell Press, 44(3), pp. 547–555. doi: 10.1016/j.neuron.2004.10.014.

Pinheiro, J. et al. (2019) ‘nlme: Linear and Nonlinear Mixed Effects Models’. R package version 3.1-141. Available at: https://cran.r-project.org/package=nlme.

Pleisch, G. et al. (2019) ‘Emerging neural specialization of the ventral occipitotemporal cortex to characters through phonological association learning in preschool children’, NeuroImage. Academic Press, 189, pp. 813–831. doi: 10.1016/j.neuroimage.2019.01.046.

Price, C. J. and Devlin, J. T. (2011) ‘The interactive account of ventral occipitotemporal contributions to reading.’, Trends in cognitive sciences. Elsevier Ltd, 15(6), pp. 246–53. doi: 10.1016/j.tics.2011.04.001.

Reynolds, C. R. and Kamphaus, R. W. (2003) ‘Reynolds Intellectual Assessment Scales: Professional manual’. Lutz, FL: PAR.

Rosenke, M. et al. (2021) ‘A Probabilistic Functional Atlas of Human Occipito-Temporal Visual Cortex’, Cerebral Cortex. Oxford Academic, 31(1), pp. 603–619. doi: 10.1093/CERCOR/BHAA246.

Sasanguie, D. et al. (2013) ‘Approximate number sense, symbolic number processing, or number-space mappings: What underlies mathematics achievement?’, Journal of Experimental Child Psychology. Academic Press, 114(3), pp. 418–431. doi: 10.1016/j.jecp.2012.10.012.

Shum, J. et al. (2013) ‘A brain area for visual numerals’, Journal of Neuroscience. Society for Neuroscience, 33(16), pp. 6709–6715. doi: 10.1523/JNEUROSCI.4558-12.2013.

Skagenholt, M., Skagerlund, K. and Träff, U. (2021) ‘Neurodevelopmental differences in child and adult number processing: An fMRI-based validation of the triple code model’, Developmental Cognitive Neuroscience. Elsevier, 48, p. 100933. doi: 10.1016/J.DCN.2021.100933.

Sokolowski, H. M. et al. (2017) ‘Common and distinct brain regions in both parietal and frontal cortex support symbolic and nonsymbolic number processing in humans: A functional neuroimaging meta-analysis’, NeuroImage. Academic Press Inc., 146, pp. 376–394. doi: 10.1016/j.neuroimage.2016.10.028.

Starrfelt, R. and Behrmann, M. (2011) ‘Number reading in pure alexia-A review’, Neuropsychologia. Pergamon, pp. 2283–2298. doi: 10.1016/j.neuropsychologia.2011.04.028.

Stevens, C. et al. (2013) ‘Relative laterality of the N170 to single letter stimuli is predicted by a concurrent neural index of implicit processing of letter names.’, Neuropsychologia. Elsevier, 51(4), pp. 667–74. doi: 10.1016/j.neuropsychologia.2012.12.009.

Stevens, W. D. et al. (2017) ‘Privileged functional connectivity between the visual word form area and the language system’, Journal of Neuroscience. Society for Neuroscience, 37(21), pp. 5288–5297. doi: 10.1523/JNEUROSCI.0138-17.2017.

Strik, W. K. et al. (1998) ‘Three-dimensional tomography of event-related potentials during response inhibition: Evidence for phasic frontal lobe activation’, Electroencephalography and Clinical Neurophysiology - Evoked Potentials. Elsevier, 108(4), pp. 406–413. doi: 10.1016/S0168-5597(98)00021-5.

Tanaka, J. W. and Curran, T. (2001) ‘A neural basis for expert object recognition.’, Psychological science, 12(1), pp. 43–7. Available at: http://www.ncbi.nlm.nih.gov/pubmed/11294227.

Tarkiainen, A. et al. (1999) ‘Dynamics of letter string perception in the human occipitotemporal cortex.’, Brain : a journal of neurology, 122 (Pt 1, pp. 2119–32. Available at: http://www.ncbi.nlm.nih.gov/pubmed/10545397.

Vogel, E. K. and Luck, S. J. (2000) ‘The visual N1 component as an index of a discrimination process.’, Psychophysiology, 37(2), pp. 190–203. Available at: http://www.ncbi.nlm.nih.gov/pubmed/10731769.

Wang, F. et al. (2020) ‘Development of Print-Speech Integration in the Brain of Beginning Readers With Varying Reading Skills’, Frontiers in Human Neuroscience. Frontiers Media S.A., 14, p. 289. doi: 10.3389/fnhum.2020.00289.

Weiss, R. H. and Osterland, J. (2013) CFT 1-R Grundintelligenztest Skala 1 - Revision. Göttingen: Hogrefe.

Wenger, E. et al. (2017) ‘Expansion and Renormalization of Human Brain Structure During Skill Acquisition’, Trends in Cognitive Sciences. Elsevier Ltd, pp. 930–939. doi: 10.1016/j.tics.2017.09.008.

Wong, A. C. N. et al. (2005) ‘An early electrophysiological response associated with expertise in letter perception.’, Cognitive, affective & behavioral neuroscience, 5(3), pp. 306–18. Available at: http://www.ncbi.nlm.nih.gov/pubmed/16396092 (Accessed: 6 March 2013).

Yeatman, J. D., Rauschecker, A. M. and Wandell, B. a (2013) ‘Anatomy of the visual word form area: adjacent cortical circuits and long-range white matter connections.’, Brain and language. Elsevier Inc., 125(2), pp. 146–55. doi: 10.1016/j.bandl.2012.04.010.

Yeo, D. J., Wilkey, E. D. and Price, G. R. (2017) ‘The search for the number form area: A functional neuroimaging meta-analysis’, Neuroscience and Biobehavioral Reviews. Elsevier Ltd, pp. 145–160. doi: 10.1016/j.neubiorev.2017.04.027.

Yoncheva, Y. N. et al. (2010) ‘Attentional focus during learning impacts N170 ERP responses to an artificial script.’, Developmental neuropsychology, 35(4), pp. 423–445. doi: 10.1080/87565641.2010.480918.

