## Appendix for "Visual occipito-temporal sensitivity to digits through elementary school"

**Supplementary Informatiom**

### 8 Appendix A: supplementary methods

#### 8.1 Overview of data available

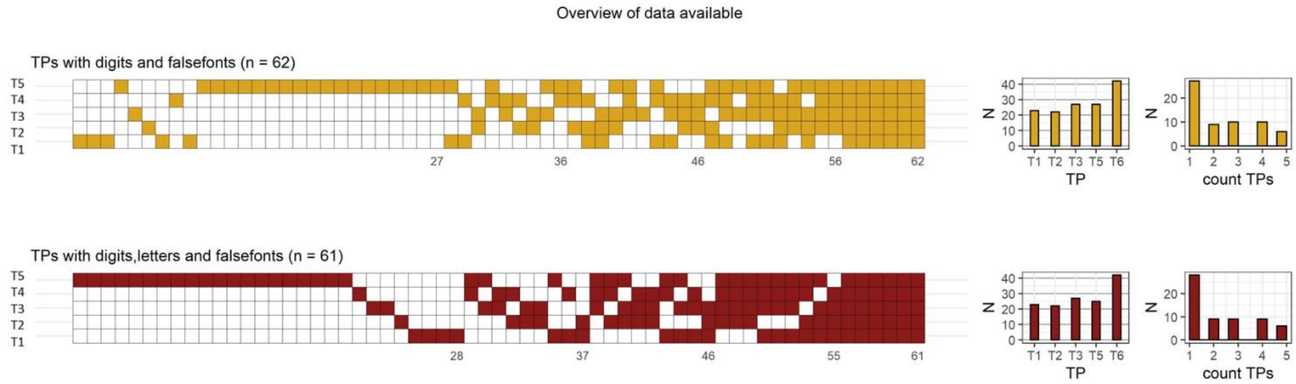


**Fig. A.1**. The tile plots on the left show longitudinal data availability for the contrasts digits vs false fonts, and for all three conditions. Columns represent participants and rows measurement time point (TP). Colored tiles indicate cases available for analysis and the x-axis labels indicate sample sizes. The bar plots summarize the sample size (N) per TP and the N depending of the number of measurements available (middle and right column of plots, respectively).

#### 8.2 Task design

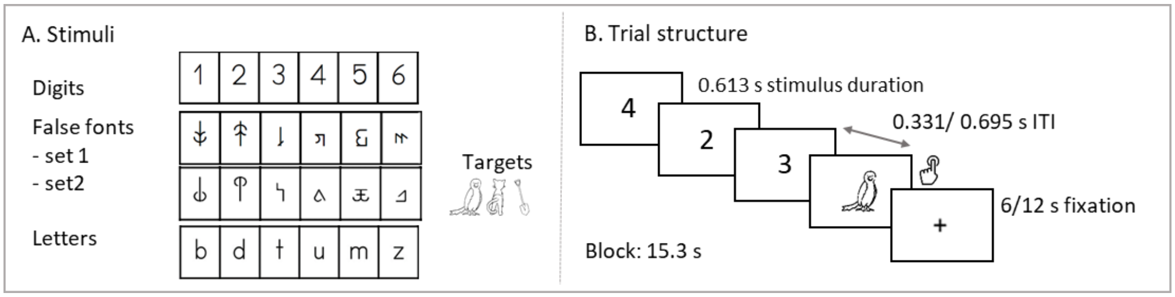


Fig. A.2. Target detection task (A) digits, false fonts, letters and target stimuli. (B) Trial structure and presentation times.

##### 8.3 Cognitive assessments

Phonological processing was assessed with subtests of two standardized tests (some subtests were not assessed after T3 as they were designed for children up to first grade). We used the rhyme and initial sound categorization subtests from TEPHOBE (Mayer, 2011). In the rhyme subtest, children are asked to select two rhyming words out of four orally presented words, whereas in the initial sound categorization subtest they are asked to select two out of four words starting with the same speech sound. Both subtests consist of seven trials. In addition, we used several subtests from the Basiskompetenzen für Lese-Rechtschreibleistungen (BAKO; Stock, Marx, & Schneider, 2013). In the phoneme deletion subtest, the children have to detect and delete the initial speech sound of a word or pseudoword and pronounce it aloud without that sound (max. seven trials). In pseudoword segmentation, children have to segment auditorily presented pseudowords by vocalizing each phoneme separately while moving tokens representing each phoneme towards the experimenter (max. eight trials). In the vowel substitution subtest, children have to repeat words while replacing all instances of the vowel [a] with [ɪ] (12 trials). The number of correct trials were used as raw scores in all subtests from BAKO test.

In addition, we assessed the rapid automatized naming (RAN) subtests of letters, numbers, colors and objects of the "Test zur Erfassung der phonologischen Bewusstheit und der Benennungsgeschwindigkeit" (TEPHOBE; Mayer, 2011).

### 9 Appendix B: Supplementary results

##### 9.1 Behavioral assessments

RAN skills showed improvements over time. RAN objects and colors were tested from T1 to T5, and RAN letters and digits from T2 to T5. The linear mixed model analysis showed a significant effect of time for RAN colors ($F$ (4,74) = 49.68, $p < 0.001$), RAN letters ($F$ (3,58) = 101.86, $p < 0.001$), RAN numbers ($F$ (3,58) = 93.42, $p < 0.001$) and RAN objects ($F$ (4,75) = 105.03, $p < 0.001$). The pairwise comparisons for RAN colors showed significant differences in T1-T2 ($t$ (74) = -3.45, $p$ = 0.008) and T4-T5 ($t$ (74) = -6.19,

$p < 0.001$), and a trend in T3-T4 ($t$ (74) = -2.72, $p = 0.060$), while the difference T2-T3 was not statistically significant, $p = 0.710$. For RAN letters, the analysis yielded significant differences in T2-T3 ($t$ (58) = -3.92, $p = 0.001$) and T4-T5 ($t$ (58) = -11.89, $p < 0.001$) but not in T3-T4, $p = 0.439$. For RAN numbers we found statistically significant differences in T3-T4 ($t$ (58) = -3.33, $p = 0.008$) and T4-T5 ($t$ (58) = -10.63, $p < 0.001$), but not in T2-T3, $p = 0.175$. Lastly, the comparisons for RAN objects yielded statistically significant differences in T3-T4 ($t$ (75) = -4.61, $p < 0.001$) and T4-T5 ($t$ (75) = -9.82, $p < 0.001$), all other *ps* > 0.129.

Regarding phonological processing assessments, we found statistically significant improvements in TEPHOBE initial sound categorization over time, $F$ (2,30) = 51.48, $p < 0.001$. The gains were significant from T1 to T2 ($t$ (30) = -7.60, $p < 0.001$), the T2-T3 comparison did not yield a significant effect, $p = 0.157$. Change with time in TEPHOBE rhyme scores was not statistically significant ($p = 0.406$). We should note that performance in TEPHOBE tests (with just 7 items) was close to ceiling levels already in T2. This was not the case for BAKO tests. The BAKO phoneme deletion test showed gains with time ($F$ (2,30) = 36.28, $p < 0.001$), that were significant in the T1-T2 comparison ($t$ (30) = -6.68, $p < 0.001$), but not in T2-T3, $p = 0.479$. For BAKO pseudoword segmentation, the main effect of time ($F$ (2,30) = 46.90, $p < 0.001$) was followed by significant effect in T1-T2 ($t$ (30) = -6.52, $p < 0.001$) and T2-T3 ($t$ (30) = -2.79, $p = 0.024$). The BAKO vowel replacement was tested in 4 timepoints (T1 to T4); statistical significance was found in the overall changes over time ($F$ *(1,* 53) = 75.81, $p < 0.001$) and the T1-T2 comparison ($t$ (53) = -10.53, $p < 0.001$), but not in T2-T3 and T3-T4 comparisons (*ps* > 0.213).

Finally, word and pseudoword reading were assessed from T2 to T5. Because different lists of items were used across time points (see section 2.2), we tested T2-T3 and T4-T5 separately and percentile scores were only compared between the latest two measurements. The analysis in the first period showed a significant increase in the number of correctly read words ($F$ (1,16) = 26.35 $p < 0.001$) and pseudowords ($F$ (1,16) = 22.23, $p < 0.001$) between T2 and T3. The comparisons of raw scores in the T4-T5 period showed significant gains for words ($F$ (1,17) = 119.18, $p < 0.001$) and pseudowords ($F$ (1,17) = 68.52, $p$ <

0.001). The analysis on percentile scores showed no statistically significant gains in words, pseudowords or the average percentile, *ps* > 0.107. Overall, the average percentile scores were relatively low performance in our sample of children at risk for dyslexia, but there was large variability between individuals. From all the children, the mean (SD; range) percentile reading scores for T4 and T5 were 26.69 (29.61; 1.25-72.75) and 32.95 (29.61; 1-96), respectively.

#### 9.2 N1 mean amplitudes for letters, digits and false fonts

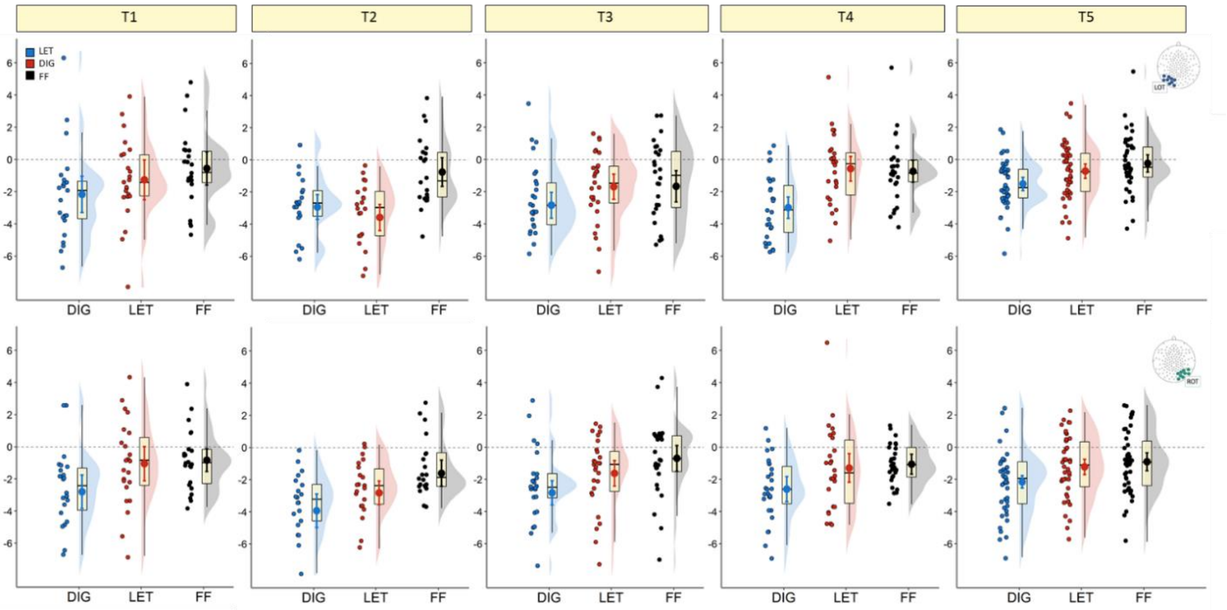


Fig.B.1. N1 mean amplitudes (μV) for digits (blue), letters (red) and false fonts (black) for left and right clusters (top and bottom row, respectively) per measurement time. Scatter points show individual means. Error bars inside boxplots show the mean and 95 % CI. LOT=left occipitotemporal; ROT = right occipito-temporal.

#### 9.3 N1 amplitude differences between digits and letters

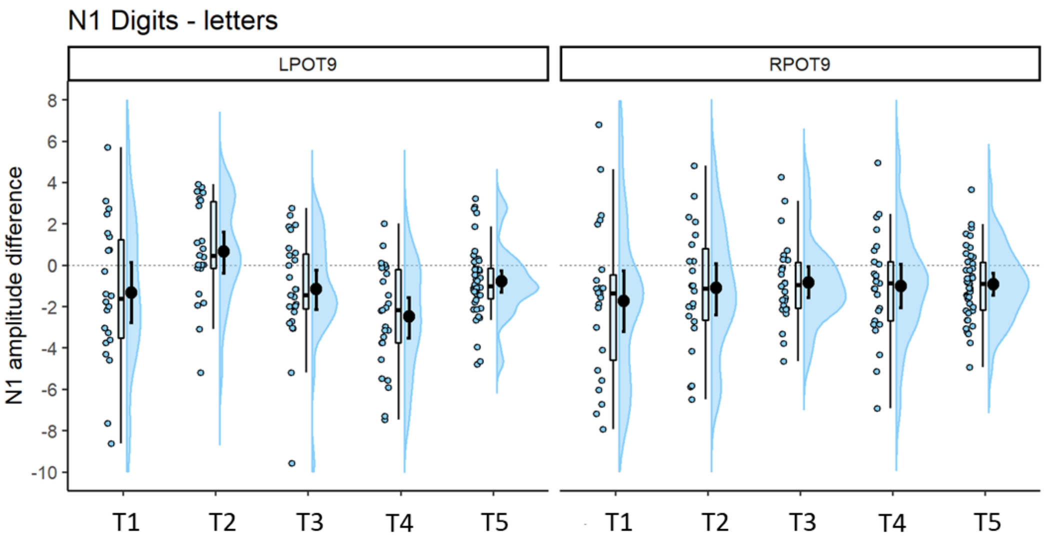


Fig.B.2. Differences in N1 amplitudes between digits and letters per time point. Error bars within the density plot indicate mean and 95 % CIs. More negative values indicate stronger negative amplitudes for digits than letters. Left hemispheric cluster is depicted on the left panel, right hemispheric cluster on the right.

#### 9.4 N1 amplitude association with arithmetic skills

The following analyses were performed separately with left and right hemisphere amplitudes. We examined the association between digit sensitivity reflected by N1 digit-false font differences in T4 and T5 and arithmetic skills in those test times. The analysis yielded a significant association between N1 digit-false font differences in the left hemisphere and the HRT scores for addition ($R$ = 0.62, $R^2$ = 0.38, β = 1.00, $t$ = 3.91, $p$ = 0.001), subtraction ($R$ = 0.44, $R^2$ = 0.19, β = 0.77, $t$ = 2.42, $p$ = 0.023), completion ($R$ = 0.53, $R^2$ = 0.28, β = 0.76, $t$ = 3.15, $p$ = 0.004), comparison ($R$ = 0.44, $R^2$ = 0.20, β = 0.87, $t$ = 2.47, $p$ = 0.021) and multiplication ($R$ = 0.51, $R^2$ = 0.26, β = 0.78, $t$ =2.97, $p$ = 0.006). There was no evidence for statistically significant associations with the HRT number writing speed, $p$ = 0.503. Interestingly, no significant associations were found between HRT arithmetic operations and N1 digit-false font difference in the right hemisphere, $ps$ > 0.171. The analysis of T5 data yielded no significant associations between N1 digit sensitivity and arithmetic skills.
